# Super-resolution optical imaging reveals accumulation of small mitochondria in the nucleoli of mouse embryos and stem cells

**DOI:** 10.1101/2023.08.03.551824

**Authors:** Kezhou Qin, Xinyi Wu

## Abstract

The nuclear mitochondrial DNA (NUMT) is found in cancer cells, but the mitochondrial DNAs entering the nuclei in normal cells have not been captured. Here, we utilized super-resolution optical imaging to capture the phenomenon by the probe PicoGreen and found mitochondrial DNAs and mitochondria accumulated in the nucleoli by four probes and overexpressing the MRPL58-DsRed. Our results provide an new explanation for mtDNA carryover and lay the foundation for the involvement of nuclear export of nucleoli in de novo mitochondrial biogenesis in another of our unpublished articles.

## Introduction

The nuclear mitochondrial DNA (NUMT) is a kind of very common phenomenon, which is not only found in cancer cells [1-3] but also present in many species [4, 5]; It’s worth mentioning that NUMT insertions are estimated to occur at a rate of ∼5 ×10^−6^ per germ cell per generation in human [5]. Besides, the mitochondria have long been found in the nuclei of patient tumors, including the adrenal cortical carcinoma of the Syrian golden hamster [6], the malignant melanoma of the iris [7], the cardiomyocytes of patients with hypertrophic and alcoholic cardiomyopathies [8], the leukemia cell [9] and the normal cow liver capsule [10], which had been proved by the transmission electron microscopy (TEM). When the mitochondria migrate towards the nucleus and accumulate near the nuclear membrane, de novo transposition from the mitochondrion to the nucleus takes place and the NUMT accumulates [2]; however, the transposition has not been captured by the super-resolution optical imaging. So, we plan to visualize the mitochondrion-related signals in the nucleus before the NUMT insertions by using several mitochondrial probes.

## Results and Discussion

Firstly, we utilized mouse embryos as the experimental materials. The animal experiment was approved by the institutional biomedical research ethics committee of the Institute of Biophysics, Chinese Academy of Sciences (permit number: SYXK202108) and was performed in compliance with the applicable institutional guidelines for the care and use of laboratory animals. We collected the mouse morula and blastosphere and detected the signals of four kinds of the mitochondrial probes including the mitochondrial DNA probe (PicoGreen), the mitochondrial reactive oxygen species (ROS) probe (MitoSOX), the mitochondrial membrane potential probes (TMRE), and the mitochondrial probes (Mito-Tracker Red). We found that all of the strongest signals of them were in the nucleolar region (Fig. S1A and B) and the nucleolar/nucleoplasmic fluorescence ratio is higher in the morula than the blastosphere (Fig. S1C). After the imaging analysis, we think that the mitochondria in the morula and the blastosphere stages of mouse embryos are immature, so it is very difficult to observe the clear mitochondrial morphology, and it is difficult to judge the occurrence of mitochondrial transfer to the nucleus.

Secondly, we used the mitochondrial probe (Mito-Tracker Green or Red, the mitochondrial probe concentration used here was 1:10000), the ROS probe (MitoSOX) and the mitochondrial DNA probe (PicoGreen) for the living cell imaging observation in human induced-pluripotent stem cells (iPSCs). Here we performed these experiments in the two states of iPSCs including the naïve state and the primed state, which can be switched to each other by simply changing the CDM medium and the RSeT medium (Fig. 1A). Through the imaging analysis, we found that the two cell states have the same phenomenon. The PicoGreen signals tend to go from the mitochondria to the nucleus (Fig. 1B and D), and are also clustered in the nucleolus zone (Fig. 1B and E). In addition, the MitoSOX signals characterizing the mitochondrial ROS level tend to enter the nucleus and accumulate clearly in the nucleolus (Fig. 1C and F). To observe that mitochondrial DNAs enter the nuclei, we performed live cell imaging observation. The PicoGreen signals in the nucleus increase with the prolonging of time (Fig. S2A and S2B). We also found that mitochondria close to the nuclei contain mtDNAs, while those far from the nuclei do not in iPSCs 2# (Fig. S2C and S2D), almost all of mitochondria doesn’t contain mtDNAs in the mesodermal cell (Fig. S2E), and small mitochondria are distributed in the nucleus and the nucleolus of the iPSCs 2# (Fig. S2D and S2F). These results demonstrate that mtDNAs and small mitochondria can enter the nuclei and accumulate in the nucleoli.

**Fig. 1.**
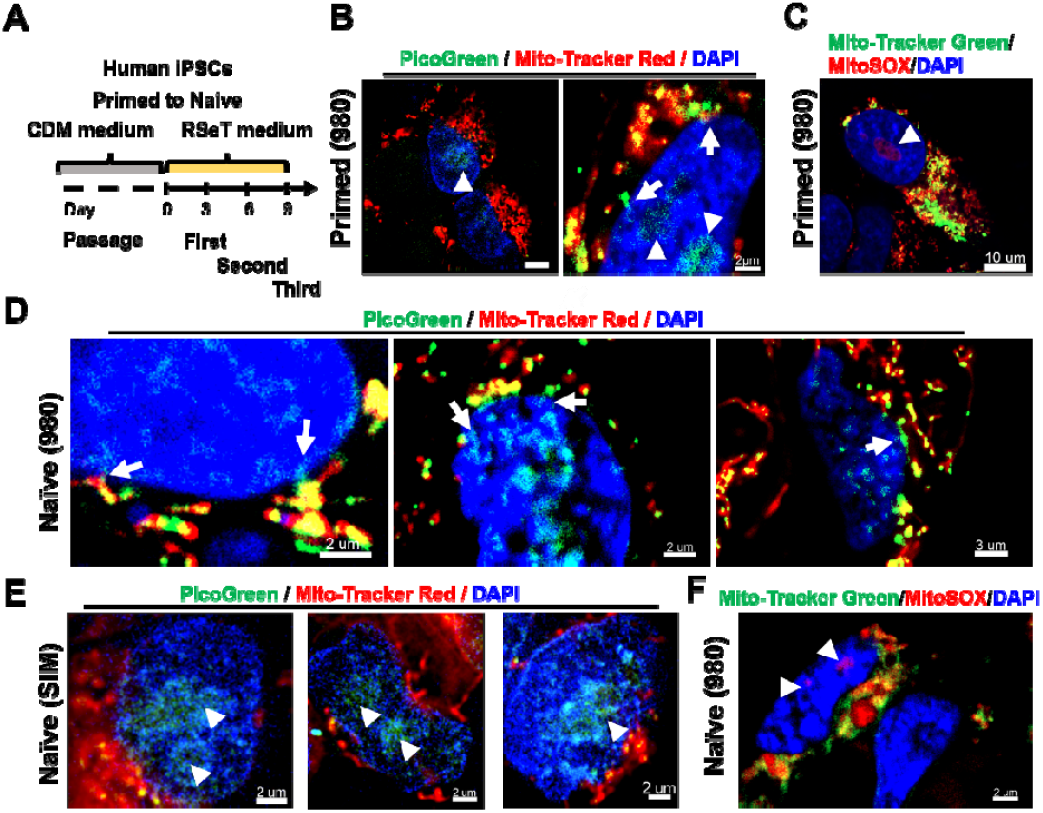
Mitochondrial probe labelling and imaging of primed and naïve human iPSCs. (A) Flow diagram of generation and maintenance of naïve human iPSCs. (B) Mitochondrial DNAs (mtDNAs) entering the nucleus (white arrows) and accumulating in the nucleoli (white arrowheads) stained with the PicoGreen and mitochondria stained with the Mito-Tracker Red in primed human iPSCs. Scale bar: 5 μm and 2 μm. (C) ROS levels in the mitochondria and a nucleolus (white arrowhead) labeled by the MitoSOX and mitochondria stained with the Mito-Tracker Green in primed iPSCs. Scale bar, 10 μm. (D) and (E) Mitochondrial DNAs (mtDNAs) entering the nucleus (white arrows) and accumulating in the nucleoli (white arrowheads) stained with the PicoGreen and mitochondria stained with the Mito-Tracker Red in naïve human iPSCs. Scale bar: 2 μm and 3 μm. (F) ROS levels in the mitochondria and a nucleolus (white arrowheads) labeled by the MitoSOX and mitochondria stained with the Mito-Tracker Green in naïve human iPSCs. Scale bars, 2 μm.

Thirdly, according to our protocol [11], we differentiated the primed human iPSCs into the mesoderm cells (Fig. 2A), and then conducted a live-cell imaging analysis, we also found the same phenomenon. Both of the PicoGreen signals and the MitoSOX signals cluster in the nucleolar region of the nucleus, and the PicoGreen signals also have a clear tendency to import into the nucleus (Fig. 2B to 2D, and Fig. S3).

**Fig. 2.**
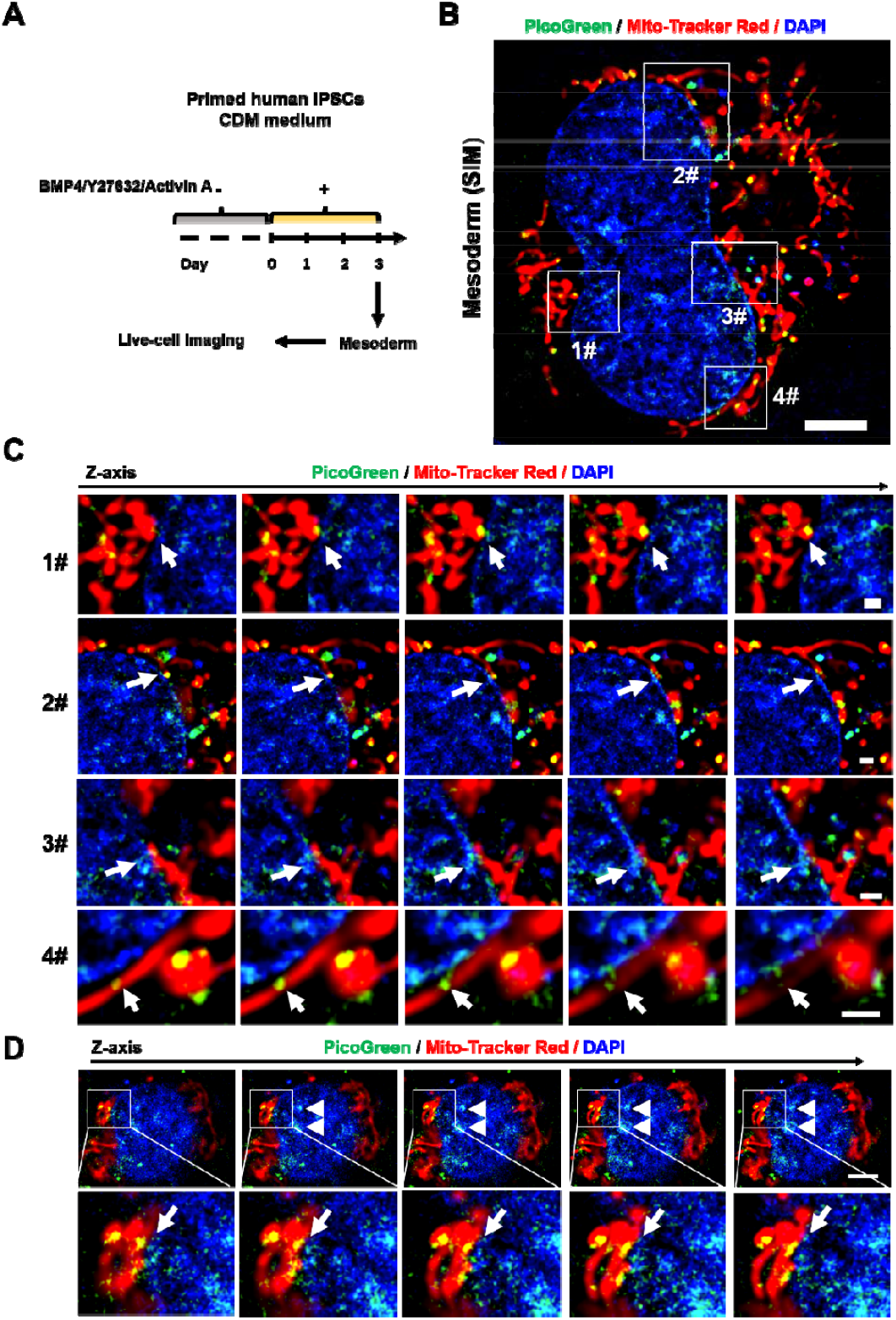
The distribution of mitochondrial DNAs (mtDNAs) in mesoderm cells stained with the PicoGreen. (A) Flow diagram of generation of mesoderm cells. (B), (C) and (D) Representative SIM images show mtDNAs are entering the nuclei (white arrows) and accumulating in the nucleoli (white arrowheads). Scale bars, 5 μm, 1 μm, 2 μm.

Fourthly, we labeled mouse embryonic stem cells (E14) with expressing the MRPL58-DsRed by using the mitochondrial probe (Mito-Tracker Green, the probe concentration is 1:1000) for the living cell imaging observation. From the above results, we confirmed that in all of these cells, the phenomenon of the mitochondria into the nucleus is present (Fig. S4). Here, we increased the working concentration of the mitochondrial probe, because we want to check whether the concentration of the probe will affect the experimental results or not. After the imaging analysis, we found that the mitochondrial probes (Mito-Tracker Green) and the mitochondrial ribosomal proteins (MRPL58-DsRed) highly overlap (Pearson’s R value: 0.87±0.05), and both of them really show that the mitochondria enter the nucleus and accumulate in the nucleolus. Taken together, the results really show us that the Mito-Tracker Green probe can mark more mitochondrial details in the nucleus by increasing the working concentration and owns a high specificity.

Summarizing the above results, we realize that there are two pathways before the occurrence of the NUMT, one is the direct entry of the mitochondrial DNAs into the nucleus, the other is the direct entry of the mitochondrial vesicles into the nucleus, and both of them finally gather in the nucleolar region (Fig. S4). In the course of these experiments, we gradually realize that the sensitivity of different mitochondrial probes is not the same, and before we perform these experiments, we may think that the specificity of the mitochondrial probe is questionable, which is because these probes can mark the nucleolar region. Now, we think that the mitochondrial probes are easier to enter the mitochondria than the nucleus, and the nucleolus, which is in the innermost layer of a cell, is the most difficult to be labeled; besides, if these mitochondrial probes are non-specific, the signals should be brighter in the nucleoplasmic region except the nucleolus.

After the PicoGreen labeling, it can be seen that there are some signals independent of the mitochondria in the cytoplasm, which is because the mitochondrial DNAs are present in the cytoplasm [12]. Although the PicoGreen labels the dsDNAs, mtDNAs are labeled brighter than the nuclear DNAs as a whole, and we can observe the phenomenon of mtDNAs entering the nuclei and accumulating in the nucleoli.

Our findings may provide a clue for mitochondrial transplantation in the treatment of mitochondrial diseases. The presence of the mitochondrial DNAs in the nucleus may affect the efficacy of mitochondrial therapy in the treatment of mitochondrial diseases. We predict that the reported mtDNA carryover [13, 14] may be related to the presence of the mitochondrial DNAs in the nucleus.

It also provides an explanation for the problem of mitochondrial contamination in the nucleolus extraction. Previously, we didn’t realize that the nucleolus might contain the mitochondria, so we have been trying to remove the mitochondrial components including the mitochondrial envelope, the mitochondrial inner membrane, the mitochondrial lumen, the mitochondrial matrix, the mitochondrial membrane, the mitochondrial nucleoid and so on [15], but in fact the nucleolus contains mitochondria. Now we correctly understand that the presence of the mitochondria-associated proteins in the nucleolus is not necessarily the result of contamination caused by the nucleolus isolation.

In conclusion, we captured the translocation of the mitochondria to the nucleus by using super-resolution microscopy and found that the mitochondria accumulate in the nucleolar region, which was confirmed by four mitochondrial probes and overexpressing one mitochondrial ribosomal protein. These results provide a new explanation for mtDNA carryover and nucleolar proteome containing mitochondrial contamination. Moreover, our results also suggest the high specificity of the mitochondrial probes and provide a new insight for the application of the mitochondrial probes. At last, this study lays the foundation for the involvement of nuclear export of nucleoli in de novo mitochondrial biogenesis in another of our unpublished articles.

## Author contributions

K.Z.Q. and X.Y.W. performed immunofluorescence assays of mouse embryos and wrote the manuscript. K.Z.Q processed all images and data analysis. K.Z.Q. directed and supervised the project.

## Acknowledgments

We thank Professor Yan Qin for support and guidance, Lei Sun for help of building mouse E14 stem cell line, for tissue section identification at the Institute of Biophysics, Chinese Academy of Sciences. We would like to thank the Center for Biological Imaging (CBI, http://cbi.ibp.ac.cn), Institute of Biophysics, Chinese Academy of Sciences for help in taking and analyzing images, and the staff from the laboratory animal research center at the Institute of Biophysics for technical assistance and efficient animal care. This work was supported by grants from the National key research and development program (2018YFC1003502 to T.J. and X.F.B.; 2018YFA0106900 to Y.Q.)

## Supplementary materials

Materials and Methods

Figures: S1 to S3

## Materials and Methods

### 1.1 Cell Generation and Maintenance

Human iPSCs 1# and 2# (given as a gift by Yue Ma lab, Institute of Biophysics in Chinese Academy of Sciences) were cultured in the hPSC-CDM™ (Cauliscell Inc. #400105) supplemented with the hPSC-CDM™ supplement (Cauliscell Inc. #600301) on the Matrigel-coated (Corning® Matrigel® Basement Membrane Matrix, *LDEV-Free, #356234) 6-well plates and were passaged with 500μM EDTA for about 3 min. Primed human iPSCs were cultured at 37°C and 5% CO2 in normal conditions. Unless otherwise specified, human iPSCs used in this article are iPSCs 1#.

Naïve stem cells were generated and stabilized in RSeT Feeder-Free Medium (Stem cell, #05975) according to the user’s guide. The naïve human iPSCs were cultured in hypoxic conditions, the medium was exchanged every other day.

Following our protocol [11], the mesoderm cells were induced from the primed human iPSCs.

Mouse E14 (given as a gift by Baoming Qin, Guangzhou Institutes of Biomedicine and Health (GIBH), Chinese Academy of Sciences (CAS), Guangzhou, China) were cultured in N2B27 based 2i+lif medium (600mL) including 288mL DMEM/F12 (HyClone), 288mL Neurobasal (Gibco), 1 % GlutaMAX (Gibco), 1 % NEAA (Gibco), 1 % Sodium Pyruvate (Gibco), 50 U/mL Penicillin/Streptomycin (Gibco), 0.1 mM β -Mercaptoethanol (Sigma), 200 x N2 (Gibco 17502001), 100 x B27 (Gibco 17504001), 1000 U/mL Lif (leukemia inhibitory factor) (Millipore), 1 μM PD0325901 (R&D, CAS 391210-10-9), 3 μM CHIR99021 (R&D, CAS 252917-06-9).

### 1.2 Animals

Animals in this study were female and male C57BL/6J mice with 4-8 weeks for only obtaining the morula and the blastosphere. All mice used were healthy and were not involved in any previous procedures. Mice were group-housed which is up to 5 animals per cage on a 12:12 hour light-dark cycle, with free access to food and water in a pathogen-free facility.

### 1.3 Collection of mice embryos at the morula and the blastosphere stage

On the first day, using 4-week-old female mice (SiPeiFu Biotech Co., Ltd.) of the C57BL/6 strain as donors, PMSG (Phenylmethylsulfonyl fluoride, Ningbo Sansheng, CasNo:329-98-6) 10 IU/mouse was injected intraperitoneally and 48 hours later, 10 IU/mouse of HCG (Human Chorionic Gonadotropin, Ningbo Sansheng, CasNo:9002-61-3) was injected intraperitoneally. Meanwhile, we took out the sperm from the 12-week-old male mice (SiPeiFu Biotech Co., Ltd.) and activated the sperms. Then, we collected the morula and the blastosphere from the developing embryos.

### 1.4 Mitochondrial staining and imaging

The mice embryos were digested with 0.1% hyaluronidase at 37°C for 15 minutes, washed three times with 1x PBS and prepared for mitochondrial staining. The prepared mouse embryos and stem cells were directly incubated with mitochondrial probes in a CO2 incubator at 37LJ and 5% CO2 or in hypoxic conditions. The cells were stained with Mito-tracker Red (100nM) (Thermal Fisher, M22425) /

Mito-tracker Green (100nM) (Thermal Fisher, M7514) for 15 min. To image mtDNAs, the cells were stained with the PicoGreen (Invitrogen, P7581) diluted 1:500 for 20 min. The Cells were incubated with 500 nM TMRE (MERK, 87917-25mg) for 10 min. The Cells were incubated with 5 μM MitoSOX (Invitrogen, M36008) for 10 min. The Cells were incubated with DAPI (MERK, Hoechst 33342, 14533-100MG) for 15 min. After incubation per time, the embryos or stem cells were washed three times with 1x PBS, cultured in the corresponding medium, and imaged with 3D using super-resolution Zeiss LSM980 (980) or structured illumination microscopy (SIM). Images were produced using an Imaris soft.

### 1.5 long-time observation of live cells

Live iPSCs 2# were cultured in confocal dishes, stained by Mito-Tracker Red, PicoGreen and Hoechst 33342, then washed three times with 1x PBS, and live□cell imaging using a live cell Imaging System is mainly performed with super-resolution microscopy Zeiss LSM980 providing fast acquisition. 3D imaging was captured with 1μm interval on Z-stack, and long-time imaging was set with 2 min intervals to visualize cell dynamics. Images were produced using an Imaris soft.

### 1.6 Statistical analysis

Statistical analyses were performed by using ImageJ and Graph Pad Prism 7. Student’s t-test was used to test for statistically significant differences and statistical significance was assumed when P <0.05.

**Fig. S1.**
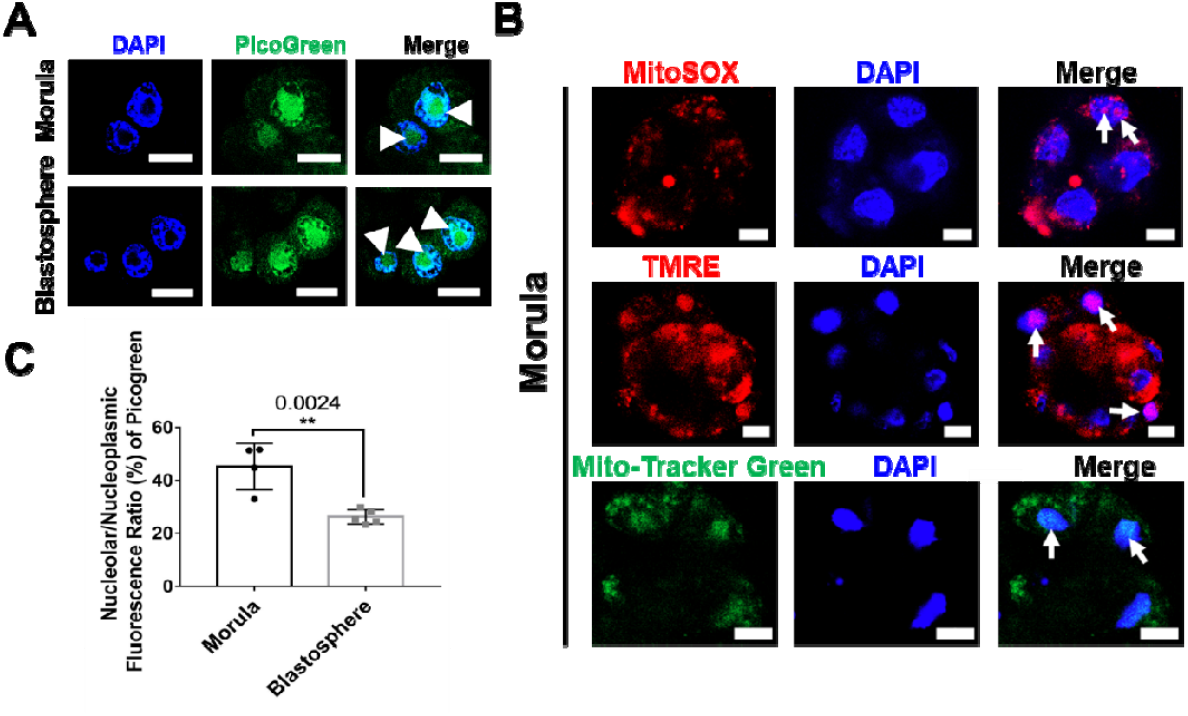
Mitochondrial probe labelling and imaging in the morula and blastosphere of mouse embryos. (A) Mitochondrial DNA content in the nucleoli (white arrowheads) stained with the PicoGreen. Scale bar, 15 μm. (B) ROS levels labeled by the MitoSOX, Mitochondrial membrane potential of Tetramethylrhodamine ethyl ester (TMRE)-stained mitochondria and mitochondria (Mito-tracker Green), which of all are checked in the nucleoli (white arrows). Scale bar, 10 μm. (C) Statistical analysis of the ratio of mitochondrial DNA content between the nucleolus and the nucleoplasm. P values were calculated using a two-tailed unpaired Student’s t-test (P values: 0.0024).

**Fig. S2.**
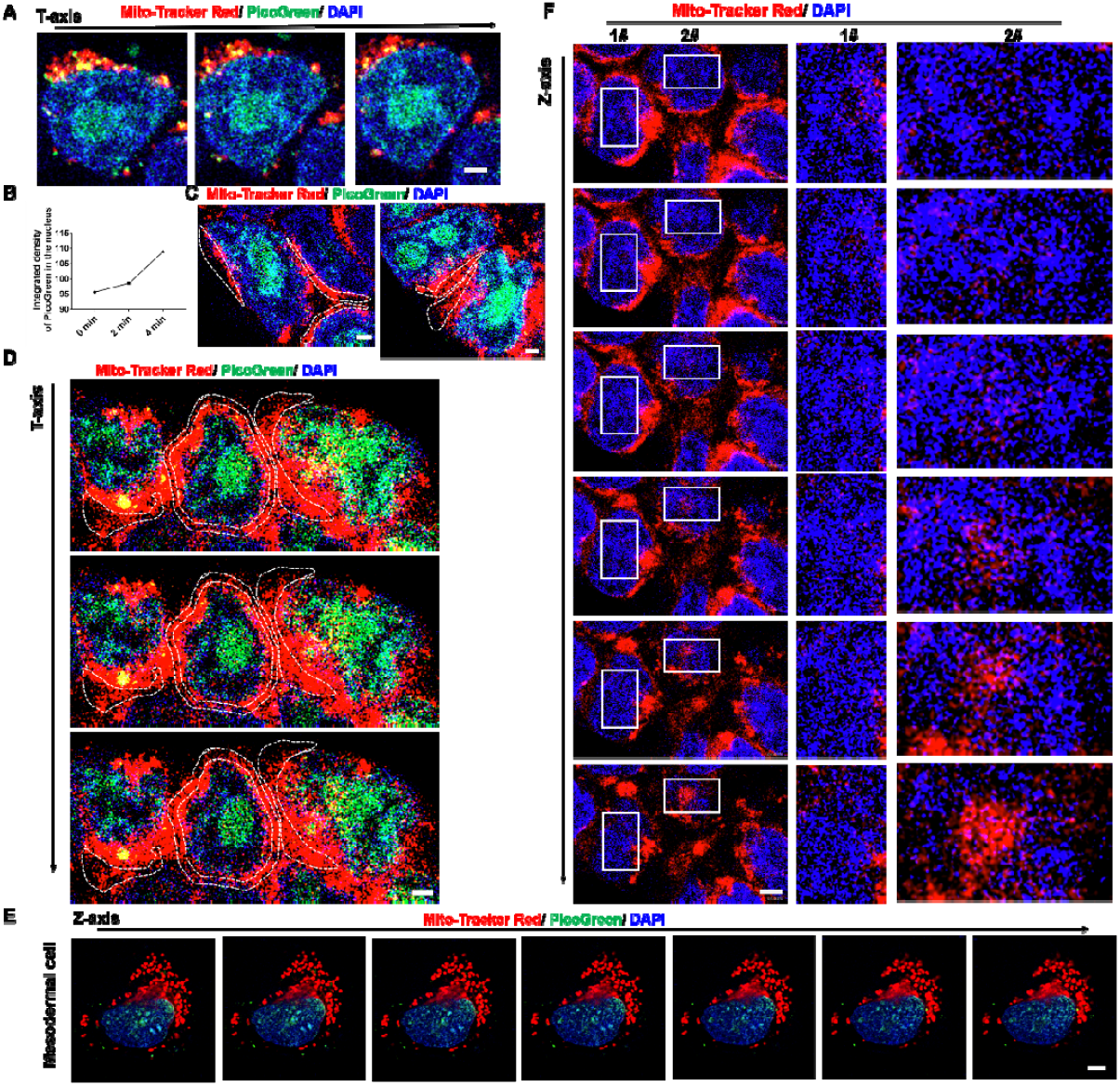
Dynamic analysis of mtDNAs, and distribution of small mitochondria in another iPSC line (iPSCs 2#) and mesodermal cells. (A) and (B) show that mtDNA content in the nuclei stained with the PicoGreen increases with the prolonging of time, and mtDNAs accumulate in the nucleolus of the iPSCs 2#. Scale bar, 2 μm. (C) and (D) show that mitochondria close to the nuclei contain mtDNAs, while those far from the nuclei (dotted boxes) do not in iPSCs 2#, and mtDNAs accumulate in the nucleoli. Scale bar, 2 μm. (E) Almost all of mitochondria doesn’t contain mtDNAs in the mesodermal cell and all mtDNAs are present in the nucleus. Scale bar, 5 μm. (D) and (F) show that distribution of small mitochondria in the nucleus or the nucleolus of the iPSCs 2#. Scale bar, 4 μm.

**Fig. S3.**
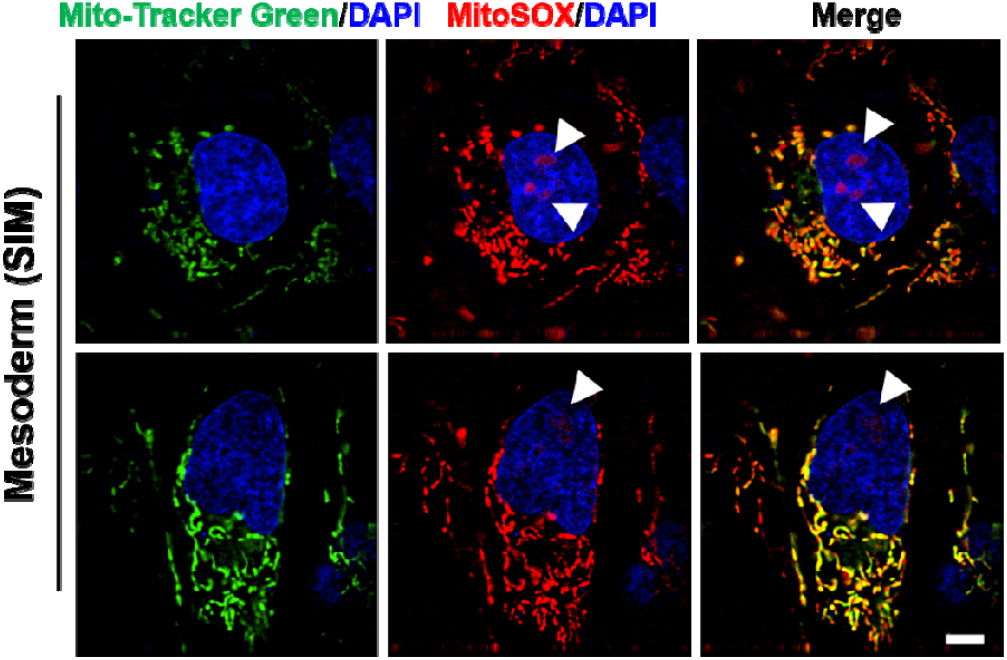
ROS levels labeled by the MitoSOX in the mesoderm cells. Representative SIM images show the ROS accumulated in the nucleoli (white arrowheads) and mitochondria stained with the Mito-Tracker Green, scale bar: 5 μm.

**Fig. S4.**
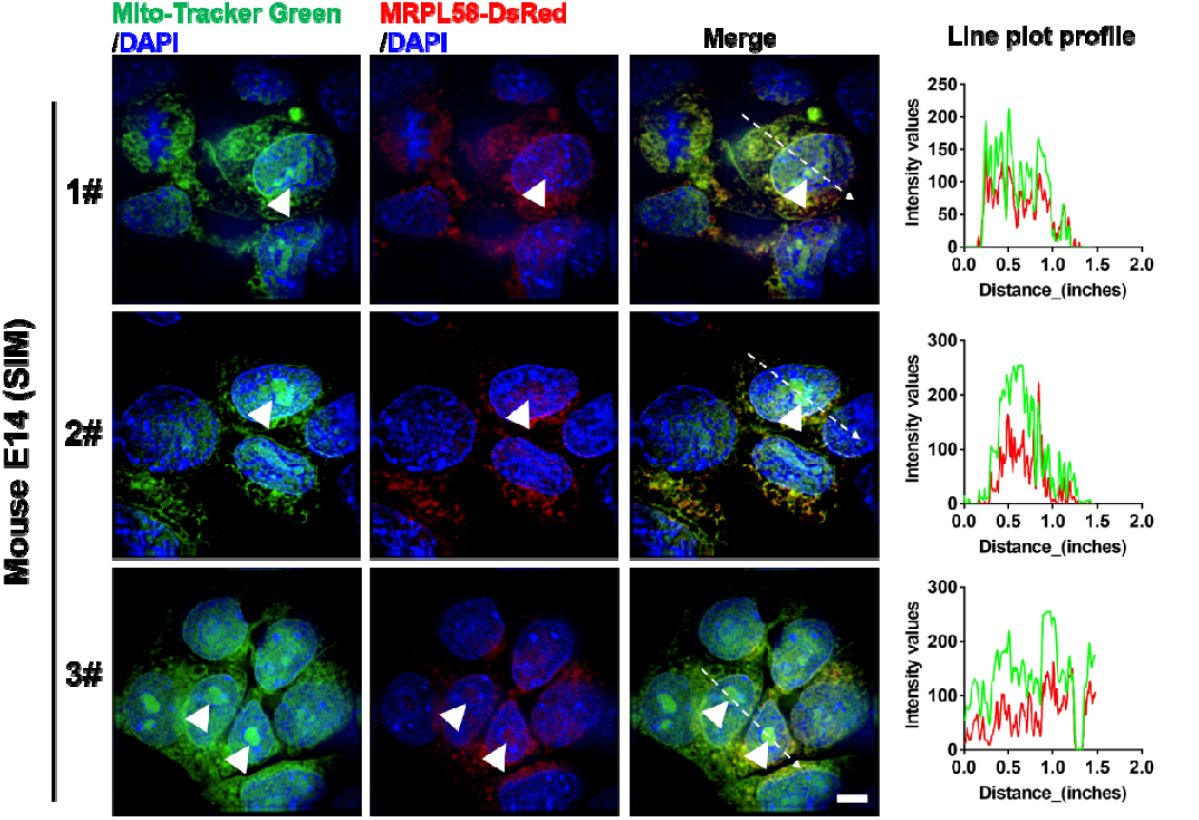
Mitochondrial probe labelling in mouse E14 stem cells with expressing the MRPL58-DsRed. Representative SIM images show colocalization of the mitochondria (Mito-Tracker Green) and the mitochondrial ribosomes, and both of them accumulated in the nucleoli (white arrowheads). Representative line profiles (dotted arrows) showed that colocalization of Mito-Tracker Green and MRPL58-DsRed (dotted arrows and blue arrow). Scale bar: 5 μm.

## References

1. Yuan, Y., Ju, Y.S., Kim, Y., Li, J., Wang, Y., Yoon, C.J., Yang, Y., Martincorena, I., Creighton, C.J., Weinstein, J.N., et al. (2020). Comprehensive molecular characterization of mitochondrial genomes in human cancers. Nat Genet 52, 342–352.

2. Srinivasainagendra, V., Sandel, M.W., Singh, B., Sundaresan, A., Mooga, V.P., Bajpai, P., Tiwari, H.K., and Singh, K.K. (2017). Migration of mitochondrial DNA in the nuclear genome of colorectal adenocarcinoma. Genome Med 9, 31.

3. Ju, Y.S., Tubio, J.M., Mifsud, W., Fu, B., Davies, H.R., Ramakrishna, M., Li, Y., Yates, L., Gundem, G., Tarpey, P.S., et al. (2015). Frequent somatic transfer of mitochondrial DNA into the nuclear genome of human cancer cells. Genome Res 25, 814–824.

4. Hazkani-Covo, E., Zeller, R.M., and Martin, W. (2010). Molecular poltergeists: mitochondrial DNA copies (numts) in sequenced nuclear genomes. PLoS Genet 6.

5. Dayama, G., Emery, S.B., Kidd, J.M., and Mills, R.E. (2014). The genomic landscape of polymorphic human nuclear mitochondrial insertions. Nucleic Acids Res 42.

6. Matsuyama, M., and Suzuki, H. (1972). Seizing mechanism and fate of intranuclear mitochondria. Experientia 28.

7. Sunba, M.S., Rahi, A.H., and Morgan, G. (1980). Tumours of the anterior uvea. II. Intranuclear cytoplasmic inclusions in malignant melanoma of the iris. Br J Ophthalmol 64.

8. Bakeeva, L.E., Skulachev, V.P., Sudarikova, Y.V., and Tsyplenkova, V.G. (2001). Mitochondria enter the nucleus (one further problem in chronic alcoholism). Biochemistry (Mosc) 66.

9. Brandes, D., Schofield, B.H., and Anton, E. (1965). Nuclear mitochondria? Science 149, 1373–1374.

10. Bloom, G.D. (1967). A nucleus with cytoplasmic features. J Cell Biol 35.

11. Qin, K., Lei, J., and Yang, J. (2022). The Differentiation of Pluripotent Stem Cells towards Endothelial Progenitor Cells - Potential Application in Pulmonary Arterial Hypertension. Int J Stem Cells 15, 122–135.

12. Shokolenko, I.N., and Alexeyev, M.F. (2015). Mitochondrial DNA: A disposable genome? Biochim Biophys Acta 1852, 1805–1809.

13. Hyslop, L.A., Blakeley, P., Craven, L., Richardson, J., Fogarty, N.M., Fragouli, E., Lamb, M., Wamaitha, S.E., Prathalingam, N., Zhang, Q., et al. (2016). Towards clinical application of pronuclear transfer to prevent mitochondrial DNA disease. Nature 534, 383–386.

14. Kang, E., Wu, J., Gutierrez, N.M., Koski, A., Tippner-Hedges, R., Agaronyan, K., Platero-Luengo, A., Martinez-Redondo, P., Ma, H., Lee, Y., et al. (2016). Mitochondrial replacement in human oocytes carrying pathogenic mitochondrial DNA mutations. Nature 540, 270–275.

15. Liang, Y.M., Wang, X., Ramalingam, R., So, K.Y., Lam, Y.W., and Li, Z.F. (2012). Novel nucleolar isolation method reveals rapid response of human nucleolar proteomes to serum stimulation. J Proteomics 77, 521–530.

